# Unexpected variation across mitochondrial gene trees and evidence for systematic error: How much gene tree variation is biological?

**DOI:** 10.1101/171413

**Authors:** Emilie J. Richards, Jeremy M. Brown, Anthony J. Barley, Rebecca A. Chong, Robert C. Thomson

## Abstract

The use of large genomic datasets in phylogenetics has highlighted extensive topological variation across genes. Much of this discordance is assumed to result from biological processes. However, variation among gene trees can also be a consequence of systematic error driven by poor model fit, and the relative importance of these biological versus methodological factors in explaining gene tree variation is a major unresolved question in phylogenetics. Using mitochondrial genomes to control for biological causes of gene tree variation, we estimate the extent of gene tree discordance driven by systematic error and employ posterior prediction to highlight the role of model fit. We find that the amount of discordance among mitochondrial gene trees is similar to the amount of discordance found in other studies that assume only biological causes of variation. This similarity suggests that the role of systematic error in generating gene tree variation is underappreciated and that critical evaluation of the fit between assumed models and the data used for inference is important for the resolution of unresolved phylogenetic questions.

---

Large genomic datasets are increasingly being used for phylogenetic inference because they increase statistical power and reduce stochastic error, which can lead to greater phylogenetic resolution (Rokas et al. 2003; Gee 2003; Rokas and Carroll 2005). The use of these large datasets has highlighted the extensive topological variation that can be found across genes. For example, phylogenomic analysis of 1,070 genes from 23 yeast genomes resulted in 1,070 distinct gene trees (Salichos and Rokas 2013). This discordance is frequently viewed as the outcome of one of several biological sources of gene variation: incomplete coalescence, horizontal gene transfer, and gene duplication/loss events in the evolutionary history of genes (reviewed by Maddison 1997; Nakhleh et al. 2013). Explicit modeling of these processes, when possible, can accommodate this variation during the inference of a species tree (Edwards 2009; Degnan and Rosenberg 2009; Boussau et al. 2013, Mirarab et al. 2014; Szöllosi et al. 2015, Edwards et al. 2016). However, variation among gene trees can also be a consequence of systematic error that arises when the model used for estimating the gene tree fits the data poorly. The relative importance of biological versus methodological factors in explaining gene tree variation is a major unresolved question in phylogenetics.

When the model fails to account for important features of the data, inferences and measures of confidence can be inaccurate (Huelsenbeck and Hillis 1993; Yang et al. 1994; Swofford et al. 2001; Huelsenbeck and Rannala 2004; Lemmon and Moriarty 2004; Brown and Lemmon 2007; Brown and Thomson 2017). Because the complexity of datasets grows with size, the potential for poor model fit to bias inferences also grows. Increasing dataset size may reduce stochastic error, but it can also exacerbate systematic error and lead to high confidence in erroneous phylogenies (Phillips et al. 2004; Delsuc et al. 2005; Jeffroy et al. 2006; Philippe et al. 2011; Kumar et al. 2012). Several cases are now known where different genomic datasets support conflicting phylogenetic hypotheses with very high statistical support (e.g. Dunn et al. 2008; Philippe et al. 2009; Schierwater et al. 2009; Whelan et al. 2015), sometimes implying very different scenarios for the evolution of important traits (e.g., the origin of nervous systems). The relative roles of biological variation and systematic error in causing this conflict is not yet well understood.

One challenge with evaluating the contributions of systematic error to gene tree discordance is that biased inferences are difficult to detect reliably given that the true evolutionary history among most taxa is unknown. However, we can greatly reduce the confounding effects of biological processes on our ability to identify systematic error by making use of the mitochondrial genome as a model system. The entire mitochondrial genome is expected to have the same evolutionary history because it is haploid and uniparentally inherited, so recombination does not typically occur. While recombination and biparental inheritance have been documented in animal mitochondrial genomes, these occurrences appear to be rare relative to the ubiquity of such events in nuclear genomes (reviewed in White et al. 2008). Therefore, analyses using individual mitochondrial genes should result in concordant gene trees. Conflict among topologies arising from different mitochondrial genes would therefore most easily be explained by systematic error during inference of gene trees.

While biased inferences are often difficult to identify directly, several approaches have been proposed to detect poor fit between models and data (e.g. Goldman 1993; Huelsenbeck et al. 2001; Bollback 2002; Nielsen 2002; Foster 2004; Rodrique et al. 2009; Ripplinger and Sullivan 2010; Brown 2014; Reid et al. 2014; Slater and Pennell 2014; Doyle et al. 2015; Duchêne et al. 2015; Barley and Thomson 2016; Gruenstaeudl et al. 2016; Duchêne et al. 2017). When fit is poor, the potential exists for inferences to be biased. However, not all instances of poor fit will result in erroneous phylogenetic estimates. Comparison of inferred gene trees and measures of model fit across tightly linked mitochondrial genes offers a unique opportunity to understand how the outcome of model fit tests relate to gene tree variation driven by systematic error. One natural approach to conducting such tests in a Bayesian framework is known as posterior prediction, wherein samples from a posterior distribution are drawn and used to simulate many replicated ‘predictive’ datasets. By comparing the predictive to the empirical datasets in various ways, the extent to which the model captures salient features of the data can be studied.

Here we analyze mitochondrial genomes for a large set of tetrapod species to characterize the extent of gene tree discordance and, using posterior prediction, begin to explore how model fit may contribute to this discord. We find that the amount of discordance among mitochondrial gene trees is similar to the amount of discordance found in studies of nuclear gene tree variation where such discordance is assumed to result from biological factors. We were able to detect systematic error related to discordance among the gene trees in this study using posterior predictive assessments. However, more work is needed to determine specific causes of poor model fit and how these drive systematic error.

## METHODS

### Datasets

We obtained all available (as of July 31^st^ 2014) whole tetrapod mitochondrial genome sequences from GenBank, which we organized into six datasets comprising the major lineages within the clade: Crocodilians (n=20), Turtles (n=53), Squamates (n=120), Amphibians (n=157), Birds (n=253), and Mammals (n=575). We extracted all 13 protein-coding genes from each mitochondrial genome based on GenBank genome annotations. Multiple sequence alignments were then constructed based on translated codons for each mitochondrial protein-coding gene in each dataset using the MUSCLE algorithm implemented in Geneious v 8 (Edgar 2004; Kearse et al. 2012).

### Initial phylogenetic analyses

For the initial phylogenetic analysis of each of the 78 gene alignments (i.e., 6 clades × 13 genes), we selected a best-fitting substitution model according to the Akaike Information Criterion (Akaike 1974) corrected for small sample size (AICc) implemented in jModelTest v 2.2 (Darriba et al. 2012). Details on the specific model chosen for each gene alignment and alignment lengths are provided in Table S1. We first obtained posterior distributions of trees and other parameters for each alignment using Markov chain Monte Carlo (MCMC) as implemented in MrBayes v 3.2.5, with the selected model and default prior settings (Ronquist et al. 2012). For each analysis, we used two independent runs (each with four Metropolis-coupled chains) and saved the state of the chains every 1000 generations. The MCMC was run until the postburn-in posterior distributions for each analysis contained 10,000 converged samples. We checked for convergence of the continuous parameters using Tracer v 1.6 (Rambaut et al. 2014) and considered a run converged when traces for all parameters appeared to be sampling from a stationary distribution and had ESS values above 1000. We assessed convergence of the tree topology using the R package rWTYv 0.1 (Warren et al. 2017). Runs were considered converged when the bipartition posterior probabilities in the MCMC chain reached a stationary frequency in the cumulative plots and showed strong correlations (Pearson’s r > 0.9) across runs.

### Characterization of gene tree heterogeneity

To characterize the extent of gene tree heterogeneity among the thirteen genes for a given clade, we calculated three different types of summary trees (majority-rule consensus tree, 95% consensus tree, and maximum clade credibility tree) from the posterior distribution for each gene and then calculated the number of incompatible splits among these gene tree estimates. We then calculated the number of incompatible splits between each pair of gene trees for a given clade (Doyle et al. 2015; available from https://github.com/vinsondoyle/treeProcessing). This measure is related to the more widely used Robinson-Fould (RF) distance (Robinson and Foulds 1981), but focuses on incompatibilities rather than bipartitions that are present in one tree but not the other. The practical effect of this change is that polytomies do not contribute to the distance. Because we are primarily interested in identifying strongly supported differences among gene trees, this was a useful property for our study. The distributions of pairwise tree-to-tree distances among genes were then visualized with violin plots using the R package ggplot2 v2.1.0 (Wickham 2009). Since we were interested in distinguishing differences among gene trees that were strongly supported (and are more likely to be driven by systematic error) from those that had little statistical support (and may simply arise from stochastic error), we focused on discordance between 95% consensus trees (calculated using Dendropy v 4.0.3; Sukumaran and Holder 2010) for the rest of the analyses in this study.

We also visually assessed gene tree heterogeneity by looking for non-overlapping sets of topologies among the thirteen genes in a low-dimensional projection of tree space created with non-linear dimensionality reduction (NLDR) using Treescaper v 1.0.0 (Huang et al. 2016; Wilgenbusch et al. 2017). Two-dimensional projections were created for each clade based on pairwise RF tree-to-tree distances of 3,250 trees taken from the posterior distributions of all genes (250 trees per gene) using the curvilinear component analysis (CCA) and stochastic gradient decent (SGD) optimization recommended in Wilgenbusch et al. (2017).

### Model performance assessment

We assessed the absolute fit of the selected models to their respective gene alignments by performing posterior predictive assessments with both data- and inference-based test statistics. Data-based test statistics measure some characteristic of the data itself (e.g., the frequency distribution of site patterns in the alignment or variation in base composition across taxa; Goldman 1993, Huelsenbeck et al. 2001) and inference-based test statistics measure some characteristic of the resulting inference (e.g., width of the posterior distribution of trees; Brown 2014). A list of the test-statistics used in this study and brief descriptions of what they measure are provided in Table 1.

**Table 1.**
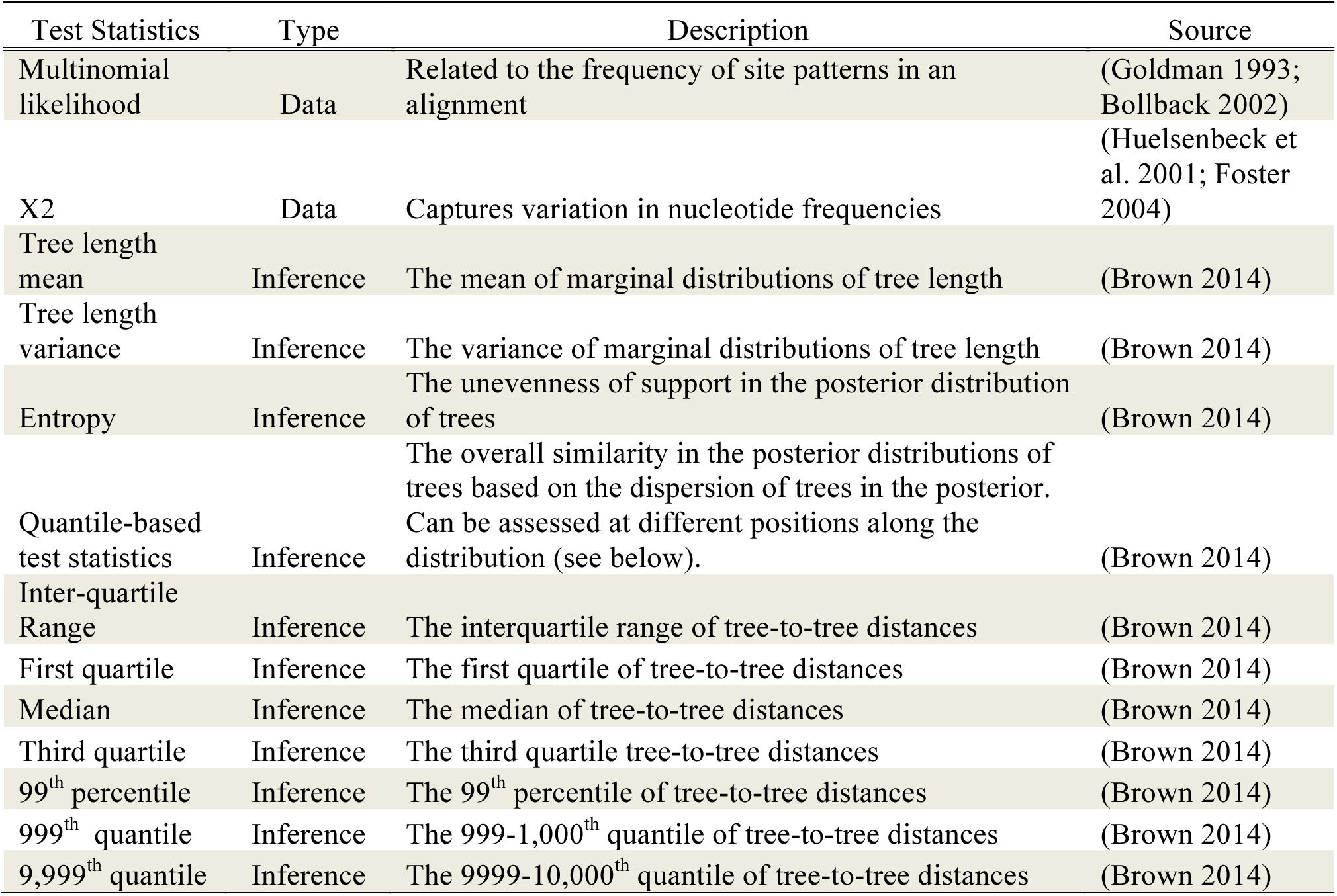
Descriptions of the model performance test statistics employed in this study. The type of test statistic refers to whether they are values based on the data themselves or the resulting inferences.

For the data-based assessments, posterior predictive simulation of datasets for each gene was performed using PuMA v0.909 (Brown and ElDabaje 2009) and SeqGen v1.3.2 (Rambaut and Grassly 1997) with 1000 parameter values and trees drawn uniformly from postburn-in MCMC samples. The data-based test statistics require that missing data be excluded from the alignments, so we removed missing data from sequences prior to simulation using PAUP^*^ v4.0b10 (Swofford 2003). Using each set of 1,000 posterior predictive datasets and the corresponding empirical dataset, we conducted two data-based assessments of model performance to characterize the ability of the model to replicate features of the empirical dataset. The multinomial likelihood test statistic (Goldman 1993; Bollback 2002; Table 1) was calculated using PuMA (Brown and ElDabaje 2009) and the *χ*^2^ statistic (Table 1) was calculated using the P4 python phylogenetics package (Foster 2004).

For the inference-based assessments, we repeated the posterior predictive simulation of datasets for each gene alignment including missing data, with 100 parameter values and trees drawn uniformly from post-burnin MCMC samples. Only 100 posterior predictive datasets were used for these tests due to the much higher computational demands involved in the inference-based assessments. For each posterior predictive dataset, we obtained a posterior distribution of trees and other parameters using MrBayes v 3.2.5 (Ronquist et al. 2012) with the model and priors assumed during analysis of the empirical data. To assess convergence, we chose five replicates at random from each gene and performed the same convergence analysis used in the initial phylogenetic analyses. When all five replicates met the convergence criteria described above, the remaining 95 predictive phylogenetic analyses were considered to have converged if the average standard deviation of split frequencies also fell below 0.01. All inference-based test statistics that were proposed in Brown (2014) were calculated in this study (Table 1) using AMP (Brown 2014, available from https://www.github.com/jembrown/amp) on a random sample of 10,000 topologies from the post-burnin posterior distribution generated for a given posterior predictive dataset. After test statistic values were calculated, we quantified the position of the empirical value relative to the posterior predictive distribution for each test using effect sizes (Doyle et al., 2015). Effect sizes for each test statistic were calculated as the absolute value of the difference between the empirical and the median posterior predictive value divided by the standard deviation of the posterior predictive distribution. These effect sizes are hereafter referred to as posterior predictive effect sizes (PPES).

### Correlation among measures of model performance

For each dataset, we ranked genes according to the model performance results and then tested for correlations among the rankings. This allowed us to assess whether the test statistics generally agreed on model performance for each gene. To do so, we calculated the rank for each gene for each test statistic based on PPES and then calculated pairwise Spearman’s rank correlation coefficients between test statistics using the R package ‘stats’ v3.2.2 (R Core Team 2015). For all pairwise combinations, we then selected one of the pair of test statistics at random and randomly shuffled its ranking of genes, recalculating the correlation coefficient. We repeated this procedure 1,000 times in order to create a null distribution of correlation coefficients and assess the significance of the observed correlation. Correlations among test statistics were considered significant if less than 5% of the coefficients from the randomized rankings were greater than or equal to the correlation coefficient from the observed rankings.

### The Relationship Between Model Fit and Gene Tree Variation

As a rough measure of accuracy in the gene tree estimates, we were interested in quantifying how different the gene trees for each clade were from widely accepted estimates of phylogeny from the literature, as well as how this might relate to measures of model performance. To do so, we selected a ‘reference tree’ from the literature for each clade that we could use as the current best estimate for that clade (Crocodilians: Oaks et al. 2011; Turtles: Thomson and Shaffer 2009; Squamates: Wiens et al. 2012; Amphibians: Pyron et al. 2011; Birds: Prum et al. 2015; Mammals: Meredith et al. 2011). Each reference tree was selected based on the availability of its posterior distributions/summary trees for analysis and similarity in taxa to those used in this study. Because we are primarily interested in strongly supported differences, we calculated the number of incompatibilities between the 95% consensus tree for each gene to the reference tree, trimming taxa as necessary so that taxon sampling matched between the two trees. We then carried out linear regression between the tree distance and the PPES for each gene and model performance test.

## RESULTS

### Agreement among gene trees from initial phylogenetic analyses

Extensive gene tree heterogeneity was present across all datasets (Fig 1). Across all datasets and consensus methods, the number of incompatibilities between genes was much greater than 0, with the exception of the Crocodilian dataset, where most genes had identical 95% consensus gene trees. The amount of disagreement varied across the types of summary tree in a way that would be expected. Maximum clade credibility trees are the most highly resolved of the summary trees, but can contain many weakly supported nodes. Thus, stochastic error in the tree estimate will increase tree-to-tree distances relative to other types of summary trees. Conversely, the 95% consensus contains fewer nodes, although all have strong support, leading to comparatively smaller tree-to-tree distances. In this latter case, the tree-to-tree distance is more likely to highlight differences that can only be explained by systematic error. Among the 95% consensus trees, tree-to-tree distances were also substantially greater than zero, indicating the presence of strongly supported yet conflicting topologies among genes. In the Crocodilian dataset containing 20 species, the majority of gene trees were well resolved and largely congruent. The conflicts among the Crocodilian gene trees occurred only among species-level relationships at the tips of the phylogenies. Gene trees for the larger datasets were less well resolved, and conflicts among gene trees in the resolution of deeper relationships were more frequent.

**Figure 1.**
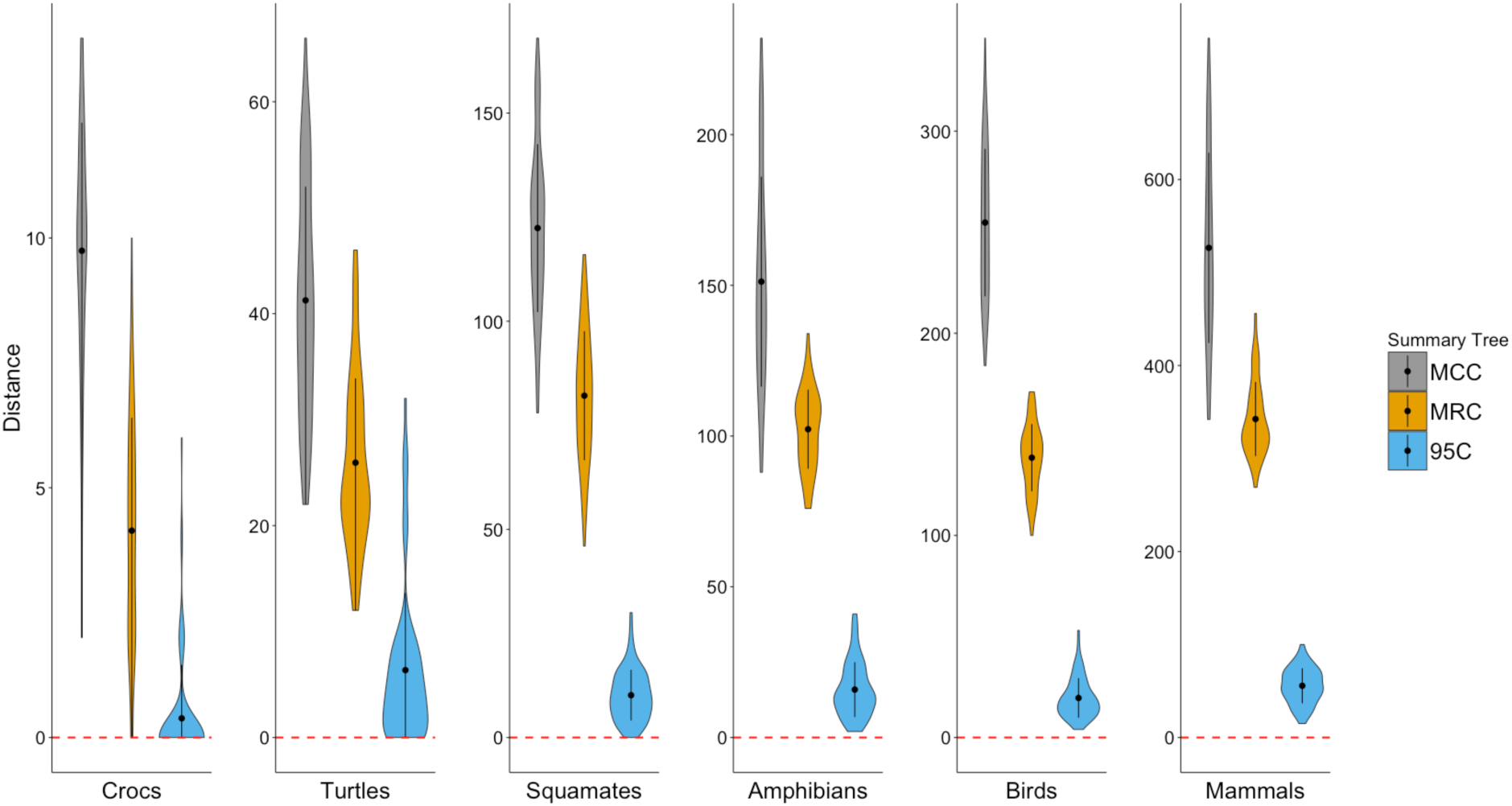
The total number of pairwise incompatibilities among all gene trees for the six datasets. Distances are shown between maximum clade credibility (MCC) trees, majority-rule consensus trees (MRC) and 95% consensus (95C) trees. The circle represents the mean number of incompatibilities and the black bars around it represent one standard deviation around the mean. The width of the violin plot indicates the density of gene trees with a particular tree-to-tree distance to another gene tree in the dataset. There is extensive variation in topology among gene trees in each clade and across summary tree types, with the exception of some 95% consensus gene trees in the Crocodilian, Turtles, and Squamates datasets.

We find similar patterns of gene tree heterogeneity in our low-dimensional projections of tree space across genes for each dataset (Fig 2). In all datasets except Crocodilians, we observe thirteen distinct clusters of trees sampled from the posterior distributions of different genes.

**Figure 2.**
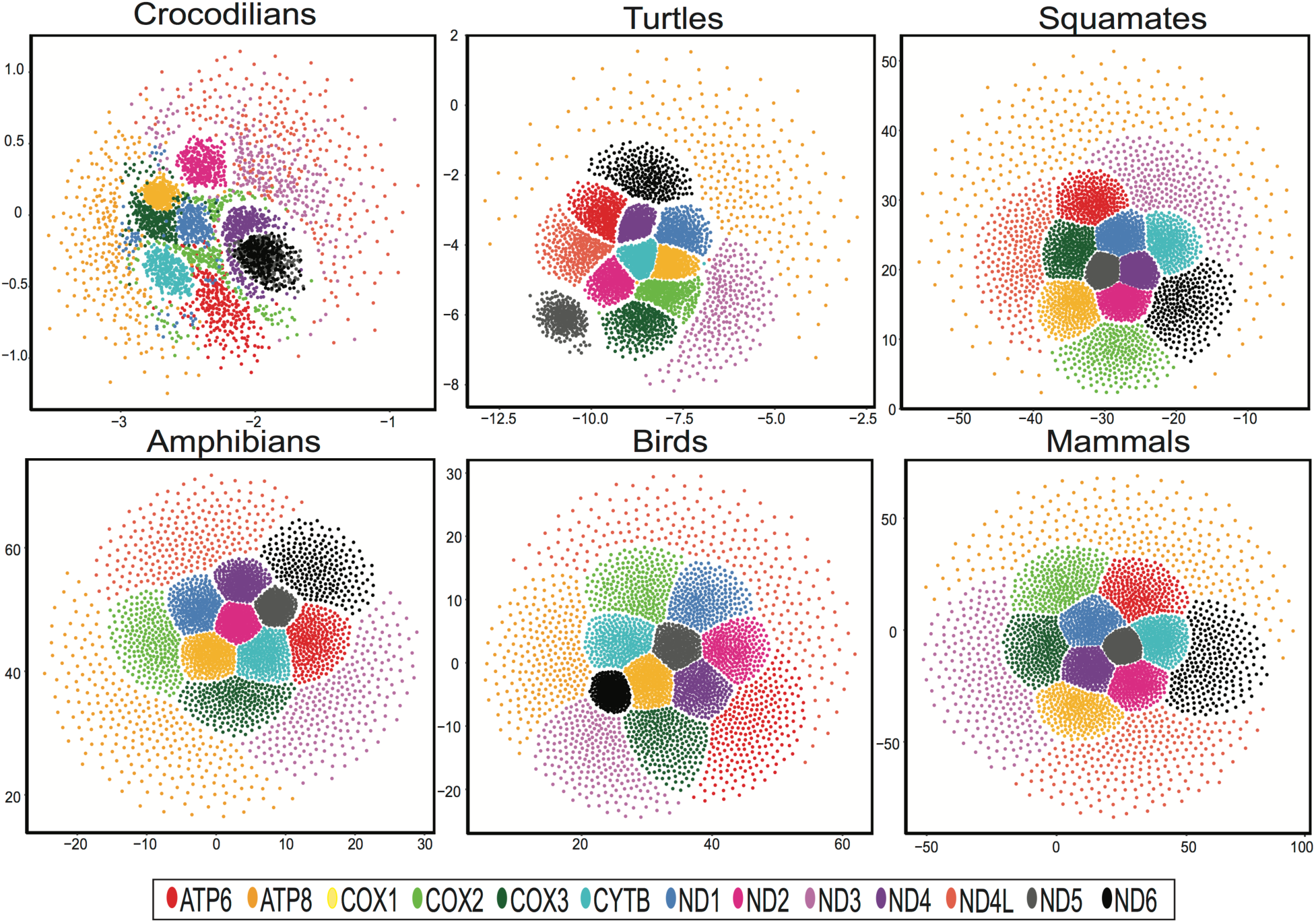
Two-dimensional NLDR representations of treespace for thirteen mitochondrial genes based on RF distances between trees. Each point represents a tree taken from the posterior distribution of a given gene.

Some of these clusters are clearly separated from other clusters (e.g. the cluster representing ND5 gene trees in the Turtle dataset), suggesting strong incongruence with other sets of gene trees.

This unanticipated level of gene tree heterogeneity across tightly linked mitochondrial genes is qualitatively similar to that found in other studies of nuclear gene tree heterogeneity (Table 2). Some of these studies (e.g. Salichos and Rokas 2013) state the observed heterogeneity could have been caused by either biological or methodological sources, and that it is nearly impossible to determine their relative contributions. Other studies (e.g. Song et al. 2012; Zhong et al. 2013; Pease et al. 2016) attribute the heterogeneity to biological factors, mainly incomplete lineage sorting, and either rule out or do not consider systematic bias as a contributing factor. Most of the above-mentioned studies characterized the extent of gene tree heterogeneity by calculating pairwise Robinson-Foulds distances among majority rule consensus trees of each locus in their dataset. We also find high levels of gene tree discordance in our mitochondrial datasets when we use similar methods for characterizing gene tree heterogeneity (Table 2), indicating that systematic bias can cause similarly extensive amounts of gene tree variation that are typically attributed to biological sources of variation.

**Table 2.** Gene tree variation found in this study compared to several other studies that focused on gene tree heterogeneity using multiple nuclear loci.

| Dataset | Taxa | Genes | Distinct Trees | Percent of possible trees found | Source |
| --- | --- | --- | --- | --- | --- |
| Crocodylians | 20 | 13 | 12 | 92 | This study |
| Turtles | 53 | 13 | 13 | 100 | This study |
| Squamates | 120 | 13 | 13 | 100 | This study |
| Amphibians | 157 | 13 | 13 | 100 | This study |
| Birds | 253 | 13 | 13 | 100 | This study |
| Mammals | 575 | 13 | 13 | 100 | This study |
| Yeast | 23 | 1070 | 1070 | 100 | Salichos and Rokas 2013 |
| Vertebrates | 18 | 1086 | 299 | 28 | Salichos and Rokas 2013 |
| Metazoans | 21 | 225 | 224 | 99.5 | Salichos and Rokas 2013 |
| Eutherian Mammals | 37 | 447 | 440 | 98.3 | Song et al. 2012 |
| Land Plants | 32 | 184 | 182 | 98.9 | Zhong et al. 2013 |
| Tomatoes | 29 | 2745 | 2743 | 99.9 | Pease et al. 2016 |

### Model Performance Assessments

The posterior predictive effect sizes resulting from the 12 model performance tests varied across genes and datasets, ranging from 0 to 1.12×10^12^ (Table 3, S2-S7). This wide range is heavily influenced by entropy, one of the inference-based test statistics, which exhibited little to no variance between posterior predictive simulations, such that small differences between the empirical and median of the posterior predictive distributions lead to extremely large PPES values for some genes in all but the Crocodilians dataset. This behavior of the test statistic stems from sensitivity to dataset size and the complexity of sampling very large tree spaces, where the coarseness of MCMC sampling makes it improbable to sample any individual topology more than once. In conventional phylogenetic analyses, where node probabilities are of primary interest, this issue is solved simply by summing up how frequently different bipartitions are sampled, rather than whole topologies. However, it becomes problematic when focusing on the frequencies of unique topologies, as we do here with the entropy test statistic. While large PPES for entropy might be meaningful for smaller datasets, it is not clear that they represent extremely poor fit between the model and the data for many of the large trees sampled here, where almost every topology sampled in the posterior is unique.

**Table 3.** The distribution of posterior predictive effect sizes (PPES) for each of the 12 model performance test statistics used in this study (Table 1) summarized across all six datasets.

| Test Statistic | Mean | St. Dev. | Median | Min | Max |
| --- | --- | --- | --- | --- | --- |
| Multinomial likelihood | 1.65 | 1.64 | 1.42 | 0.002 | 11.4 |
| X2 | 19.61 | 23.48 | 11.7 | 0.04 | 110.68 |
| Tree length mean | 1.91 | 1.85 | 1.35 | 0.026 | 8.21 |
| Tree length variance | 5.45 | 6.52 | 3.45 | 0.33 | 32.76 |
| Entropy | $6.61 \times 10^{10}$ | $2.48 \times 10^{11}$ | 0.96 | 0 | $1.12 \times 10^{12}$ |
| Interquartile range | 4.82 | 3.41 | 4.31 | 0 | 16.24 |
| 1<sup>st</sup> quartile | 4.77 | 3.02 | 4.9 | 0 | 11.73 |
| Median | 4.95 | 3.16 | 5.19 | 0 | 12.28 |
| 3<sup>rd</sup> quartile | 5.19 | 3.09 | 5.37 | 0 | 12.65 |
| 99<sup>th</sup> quantile | 5.58 | 3.29 | 5.93 | 0 | 13.44 |
| 999<sup>th</sup> quantile | 5.73 | 3.36 | 6.12 | 0 | 13.82 |
| 9999<sup>th</sup> quantile | 5.79 | 3.35 | 6.27 | 0 | 13.89 |

When entropy was excluded, data-based test statistics appeared to reject model fit among genes more strongly than inference-based test statistics across all six datasets, with larger PPES on average (Table 4). This result makes sense, since poor model fit must manifest itself at the level of the data in order for inferences to be affected, but not all model deficiencies noticeable in the data will affect inference. PPES ranged from 0.002 to 110.78, indicating a large range of model fit to the empirical data. The range of PPES for inference-based test statistics was smaller than for data-based test statistics and this range varied across datasets (Table 4). For Crocodilians, PPES across inference-based test statistics were typically small, ranging from 0 to 3.16 (Table 4 and S2), suggesting that the selected models appear to fit the Crocodilian gene alignments better than for the other datasets, although this may be due to differences in power of the test statistics to detect poor model performance across datasets of different sizes. For Turtles, PPES ranged from 0 to 14.25 (Table 4 and Table S3), indicating a mixture of model fit. Similar patterns of a mixture of model fit across genes were also found for the larger datasets of Squamates, Amphibians, Birds, and Mammals (Table 4, S4-7).

**Table 4.** The distribution of posterior predictive effect sizes (PPES) for each dataset across 11 of the 12 test statistics. Entropy was removed from the pool of test statistics summarized in this table because of the extreme outlier PPES of this test statistic across the majority of the datasets (see text). The PPES for entropy test statistic are provided in Supplementary Tables 4-9.

| Dataset | Data-based test statistics |  |  |  |  | Inference-based test statistics |  |  |  |  |
| --- | --- | --- | --- | --- | --- | --- | --- | --- | --- | --- |
|  | Mean | St. Dev. | Median | Min | Max | Mean | St. Dev. | Median | Min | Max |
| Crocs | 4.15 | 4.57 | 1.58 | 0.16 | 13.92 | 1.05 | 0.78 | 1.03 | 0 | 3.16 |
| Turtles | 3.48 | 4.02 | 1.78 | 0.04 | 15.08 | 2.21 | 2.04 | 1.97 | 0 | 14.25 |
| Squamates | 12.29 | 14.77 | 3.05 | 0.002 | 49.1 | 6.15 | 3.04 | 6.14 | 0.21 | 28.54 |
| Amphibians | 27.82 | 36.06 | 5.58 | 0.09 | 110.68 | 5.99 | 2.37 | 5.67 | 1.47 | 18.08 |
| Birds | 5.29 | 6.17 | 1.86 | 0.09 | 21.46 | 5.19 | 2.37 | 5.47 | 0.07 | 10.81 |
| Mammals | 10.77 | 13.39 | 8.63 | 0.13 | 45.89 | 8.63 | 3.99 | 9.62 | 0.87 | 16.24 |

### Correlation Among Measures of Model Performance

Across all datasets, gene rankings were significantly correlated among the quantile-based test statistics that compare the support and similarity of trees across the empirical and predicted posterior distributions (Fig 3). Within the Crocodilian and Squamate datasets, the gene rankings for the mean and variance of tree length were significantly correlated with each other. Within the Crocodilian dataset, gene rankings based on entropy were correlated with gene rankings among the quantile-based test statistics. We observed a few other correlations, although these were largely weak and idiosyncratic among datasets (Fig 3).

**Figure 3.**
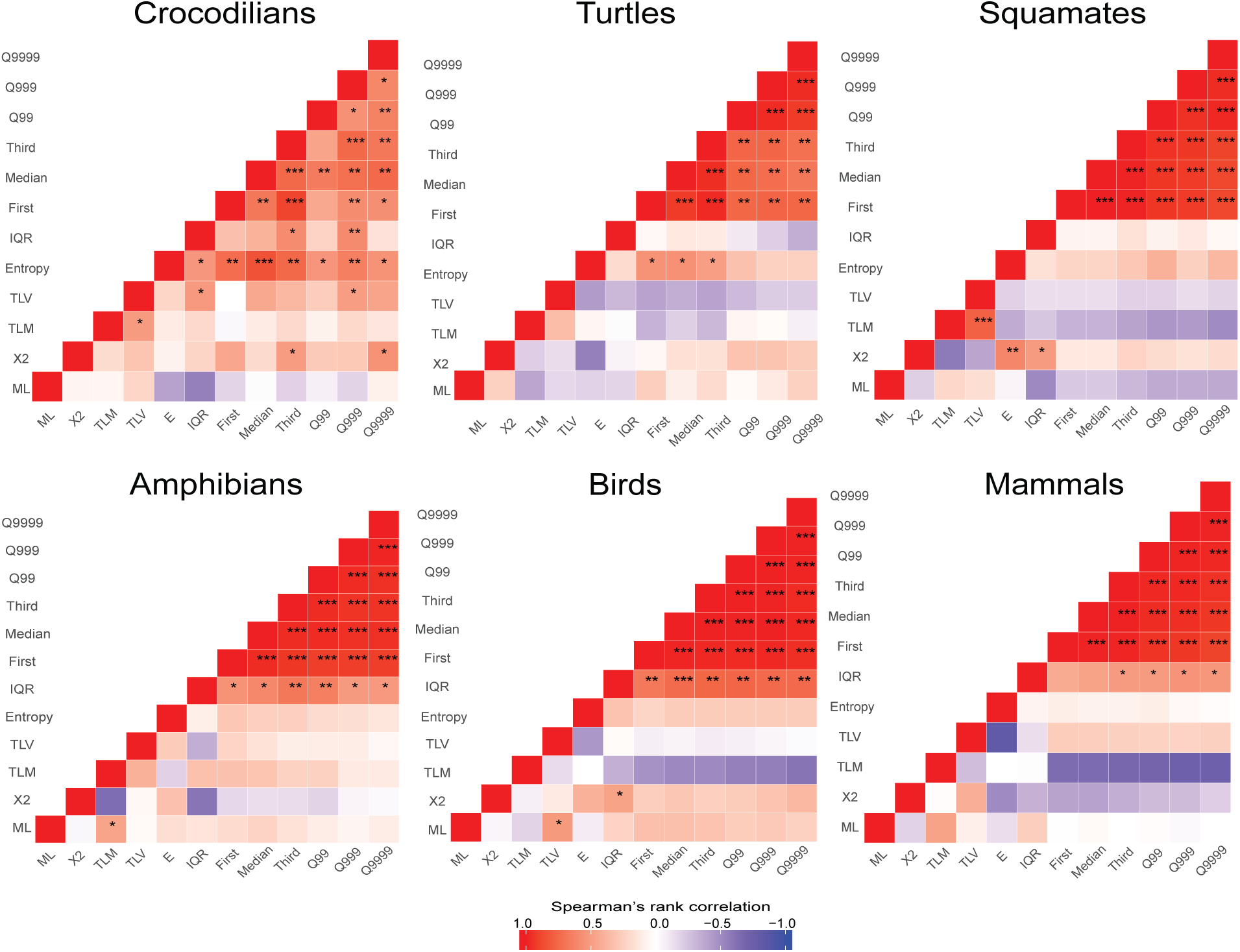
Heatmap of the Spearman’s rank correlation coefficient between gene rankings among model performance tests based on posterior predictive effect sizes. Model performance tests include multinomial likelihood (ML), composition heterogeneity (X2), tree length mean (TLM), tree length variance (TLV), statistical entropy (E), interquartile range (IQR), first quartile (First), median, third quartile (Third), 99th percentile (Q99), 999th-1000 quantile (Q999), 9999-10000th quantile (Q9999) of tree to tree distances in posterior distributions. Stars indicate correlations that are significant at a significance threshold of 0.05 (^*^), 0.01 (^**^), and 0.001(^***^).

### The Relationship Between Model Fit and Gene Tree Variation

The amount of strongly supported conflict between gene trees and reference trees varied across datasets and was low overall for Crocodilians and Birds and somewhat higher in the other clades (Table 5). There was no simple overall relationship between tree distance and PPES (Fig 4, Table S8). Although genes did vary in their PPES, increasing PPES did not necessarily correspond to decreasing congruence between gene trees and reference trees across all datasets. However, we did observe some significant positive correlations between PPES and incongruence with the reference tree (e.g. for the 999-1,000^th^ and 9999-10,000^th^ quantile-based test statistic in the Turtle dataset; Figure 4). We also observed some significant negative correlations in the same test statistics for the Crocodilian and Bird datasets. The negative relationships in these datasets may have to do with the combined effects of (1) a lack of strong disagreement among the gene trees and the reference tree (Table 5) and (2) an interaction between the power of a test statistic to detect poor model performance with the power of a gene to precisely estimate the phylogeny (i.e., the shortest genes often have the smallest PPES as well as the fewest incompatibilities with the reference tree due to lack of information rather than poor fit of the model).

**Table 5.** The percentage of bipartitions agreed upon by gene trees and reference trees for each clade. The number of taxa in each dataset after trimming is provided in parentheses. The percentage of bipartitions agreed upon was calculated the number of compatible nodes divided by the total number of nodes in the tree.

| **Gene** | **Crocs (20)** | **Turtles (49)** | **Squamates (35)** | **Amphibians (28)** | **Birds (33)** | **Mammals (104)** |
| --- | --- | --- | --- | --- | --- | --- |
| ATP6 | 95 | 88 | 77 | 96 | 97 | 88 |
| ATP8 | 95 | 98 | 94 | 96 | 97 | 99 |
| COX1 | 95 | 98 | 83 | 86 | 100 | 93 |
| COX2 | 85 | 98 | 94 | 86 | 94 | 94 |
| COX3 | 95 | 92 | 89 | 93 | 97 | 91 |
| CYTB | 90 | 92 | 97 | 100 | 100 | 83 |
| ND1 | 95 | 98 | 94 | 100 | 97 | 87 |
| ND2 | 90 | 100 | 80 | 89 | 97 | 90 |
| ND3 | 95 | 96 | 97 | 96 | 97 | 96 |
| ND4 | 90 | 90 | 94 | 96 | 100 | 89 |
| ND4L | 95 | 84 | 100 | 96 | 100 | 98 |
| ND5 | 90 | 67 | 86 | 89 | 97 | 74 |
| ND6 | 95 | 96 | 97 | 93 | 100 | 90 |

**Figure 4.**
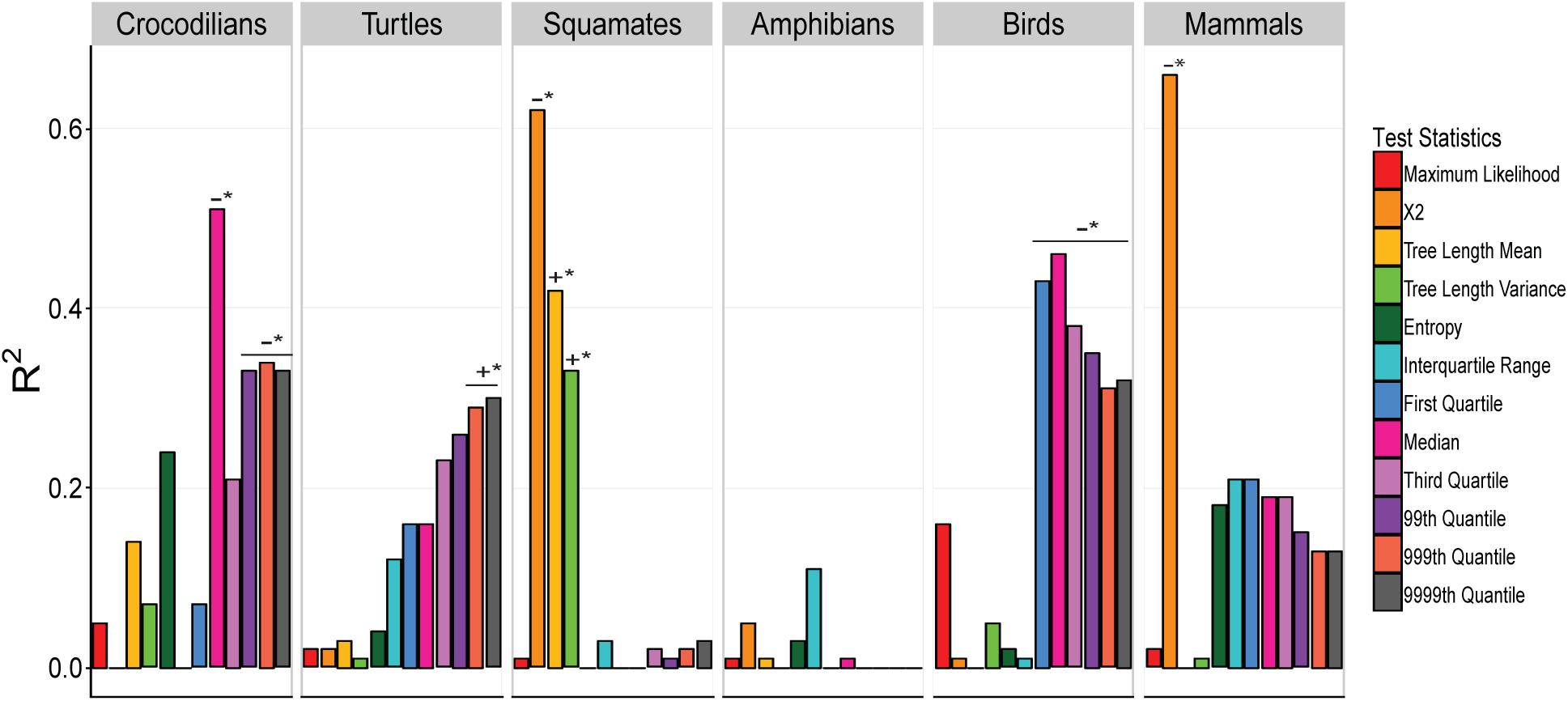
Relationship between PPES and the number of incompatibilities between 95% consensus gene tree and reference tree based on linear regression. Correlations with significantly positive or negative slopes are represented by (+^*^) and (-^*^) respectively. The values of the slope and 95% confidence intervals are provided in Supplementary Table 11.

While the relationship between poor model fit and topological conflict between the gene trees and reference tree appears to be complex, we do find several cases where these methods clearly identified systematic bias or other issues in the data. While inspecting PPES results, we noted two cases where a single gene was a large outlier for one or more model performance tests relative to all genes (Fig 5). In both cases the PPES outlier was correctly signaling an issue in the analysis. Specifically, phylogenetic analysis of cytochrome-B (CYTB) in the Squamate dataset inadvertently included a misaligned region that affected four sequences. This misalignment increased the tree length mean and variance PPES for this gene, which were consequently much larger than these values for all other genes in the dataset (Fig 5A). The error also drove a spurious phylogenetic result that united a worm lizard with several blindsnakes as a (clearly erroneous) clade. Once we corrected the misalignment, the tree length mean and variance PPES for CYTB were drastically reduced and the position of these taxa in the gene tree returned to their more commonly accepted positions.

**Figure 5.**
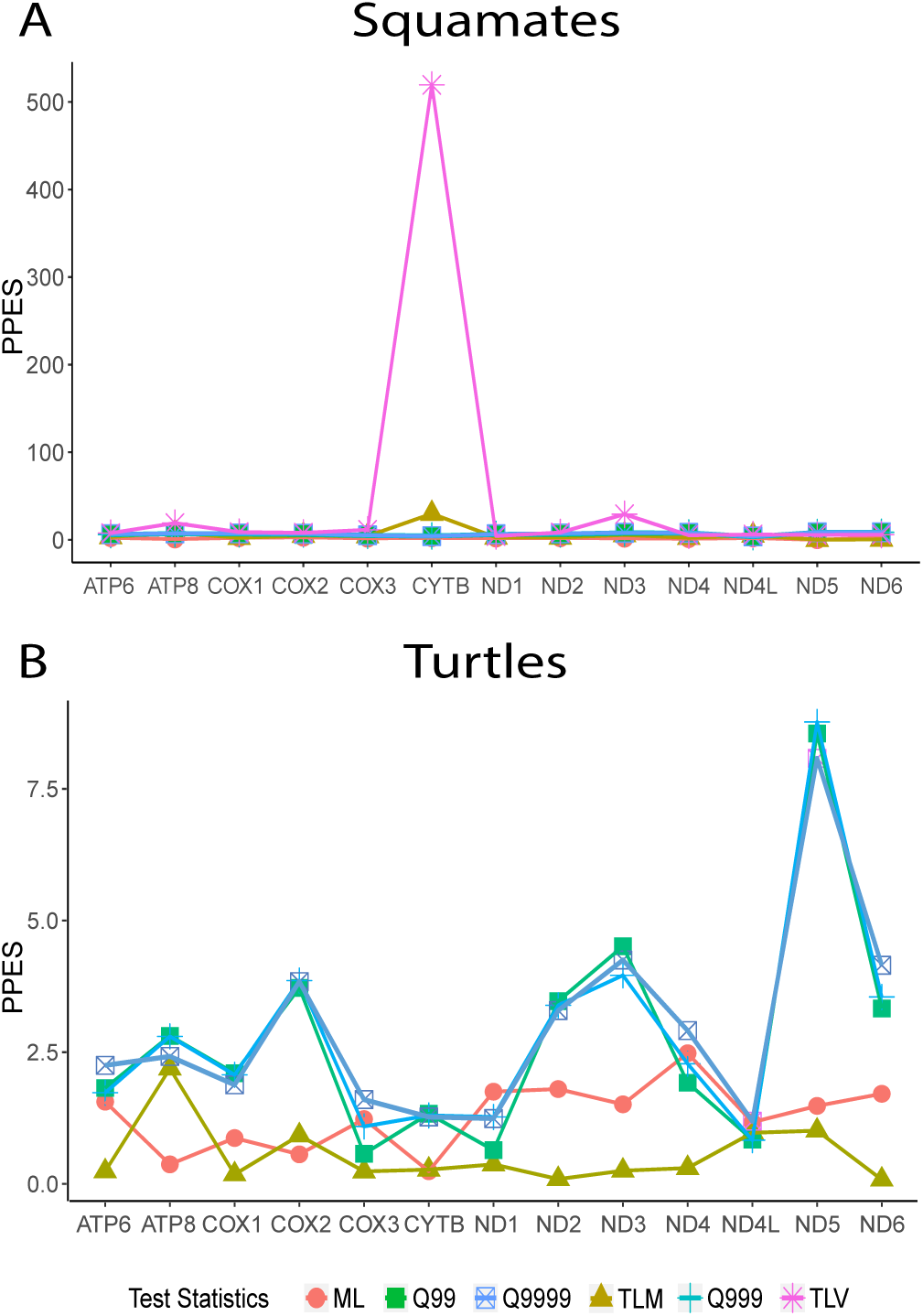
The PPES for each gene from a subset of model performance tests that highlight issues in the analysis. A) In the Squamate dataset, the PPES (before the misalignment was corrected) associated with the tree length mean and variance test for the CYTB alignment are much larger than for the other genes. B) In the Turtle dataset, the PPES associated with the quantile-based model performance tests of the ND5 alignment are twice as large as the PPES for ND3, the gene with the next largest PPES. Model performance tests shown here are the multinomial likelihood (ML), tree length mean (TLM), tree length variance (TLV), 99^th^ percentile (Q99), 999^th^-1,000 quantile (Q999), 9999-10,000^th^ quantile (Q9999) of tree-to-tree distances in posterior distributions.

The quantile-based test statistics that measure the spread of the distribution of trees within the posterior distribution also detected clear systematic error in the inference of the Turtle ND5 gene tree. The ND5 PPES for the 99-100th, 999-1000th, and 9999-10000th quantiles were at least twice as large as any other gene (Fig 5B). The gene tree for ND5 supports a fundamentally different backbone of family-level relationships among turtles and contains a large number of topological conflicts with the reference tree in comparison to the rest of the gene trees in the Turtle dataset (Table 5). Because the backbone relationships of turtles are well established (Thomson and Shaffer 2010; Barley et al. 2010; Crawford et al. 2015, Shaffer et al. 2017), we are confident that the ND5 gene tree is being influenced by systematic error. Supporting this, there was a significant positive correlation between the number of incompatibilities and model performance based on the quantile-based test statistics for this dataset (Fig 4, Table S8).

## DISCUSSION

Our analysis highlights several issues that should influence methodological choices for researchers moving forward. Most significantly, we find that the amount of gene tree variation in empirical data can be large, irrespective of whether biological sources of gene tree variation (i.e incomplete lineage sorting) are expected to play a significant role. The gene tree heterogeneity observed in this study is qualitatively similar to other studies that attribute the variation solely to biological processes. This similarity suggests that the observation of variation among gene trees in empirical data should not necessarily be ascribed to biological sources by default and researchers should take care to check for more prosaic explanations of gene tree variation in their data (i.e. poor model fit driving systematic error) before applying a hierarchical model of gene tree variation (and assuming it can adequately account for this variation). ‘Species tree’ approaches to analyzing multilocus alignments typically assume that the only source of discordance is biological (i.e. coalescent stochasticity). Other factors, such as discordance caused by poor model fit at the DNA sequence level, can contribute to gene tree heterogeneity and mislead these approaches (e.g. see Scornavacca et al. 2017 for an example where incomplete lineage sorting is only a minor cause of observed phylogenetic discordance in placental mammals). With increasing application of genomic data and the strong statistical power it provides for phylogenetic inference, it is important that researchers take into account both methodological and biological sources of gene tree conflict in the effort to produce accurate, highly supported trees.

The combination of pervasive gene tree variation coupled with the substantial evidence for systematic error suggests that, even in genomes that have been characterized and analyzed extensively (such as the mitochondrial genome), phylogenetic analyses still have the potential to be substantially mislead. In larger datasets, such as those that sample hundreds or thousands of less well characterized loci from the nuclear genome, this potential grows further. The utility of the mitochondrion for this study is that we have a strong *a priori* expectation that gene trees will be concordant in the absence of poor model fit. This expectation does not hold for larger nuclear datasets, so detecting these issues is consequently both more difficult and more critical. We attempted to use variation in model fit to sort genes into those that are more or less reliable, but found that this relationship was too complex relative to the small number of genes in the mitochondrial genome to allow for such coarse characterization. Nevertheless, this approach does appear to be fruitful when more loci are available (Doyle et al. 2015).

Model fit tests employing posterior predictive simulation, and related approaches, have the potential to fill an important gap in phylogenetic methodology by assessing a model’s fit to a given dataset. Model fit testing in a posterior predictive framework allows a great deal of flexibility to focus on different aspects of a model and their influence on inferences. In this study, we conducted a suite of model performance tests to explore possible sources of systematic error that may be driving extensive gene tree variation. Across several datasets, we were able to detect the presence of systematic error with some of the test statistics, particularly the upper quantile-based test statistics. However, the relationship between model performance and tree-to-tree distances appears to be more complex than a simple linear correlation.

This complex relationship may stem from poor performance across all genes, leading to consistent or very subtlety different levels of error across gene trees and difficulty in detecting a relationship with gene tree congruence. Alternatively, poor model performance in some genes may result in many subtle errors in estimated support for relationships in the posterior distribution that lead to large PPES values from the predictive datasets, but not result in any one part of the tree strongly conflicting with the reference (e.g. discordance among nodes deeper in the tree that cause larger tree-to-tree distances). It is also possible that the true mitochondrial history in some of these datasets, especially those that have undergone rapid radiation, may be different than the true species history.

The specific causes of poor model fit, and their role in producing systematic error, were difficult to determine with the model performance tests used here. The implementation of more site-specific and branch-specific test statistics in the posterior predictive framework could help pinpoint the specific causes of poor model fit and the regions of the tree that are most directly affected. Our difficulty with determining the sources of systematic error in this study may also stem from issues with the power of these tests to detect poor performance, as they might represent conservative measures of poor model performance (Bollback 2005, Ripplinger and Sullivan 2010, Brown 2014). The power of posterior predictive tests to detect poor model performance in a gene and the power of the gene to precisely estimate the phylogeny are probably correlated. Precise characterization of this relationship will require simulation studies beyond the scope of this paper.

## CONCLUSIONS

Gene tree heterogeneity in multilocus studies is often assumed to stem from biological processes, such as incomplete lineage sorting or horizontal transfer, and several methods have been developed to model these types of variation. We demonstrate that systematic error can be as significant a source of variation among gene trees as biological sources, although it is not currently standard practice to check for this. The posterior predictive framework for model performance assessment has the potential to fill this important gap in current phylogenetic methodology and provides researchers with a great deal of flexibility in testing different aspects of model fit. With increasing application of genomic data and the strong statistical power it provides for phylogenetic inference, it is important that researchers better take into account the methodological sources of gene tree conflict alongside the biological in the effort to produce accurate, highly supported trees.

## SUPPLEMENTARY MATERIAL

Data files and other supplementary information related to this article have been deposited at Dryad under doi:XXX

## ACKNOWLEDGMENTS

We utilized high-performance computing resources from the University of Hawai’i (UH) and Louisiana State University (LSU) for many of the analyses conducted in this study. We thank Floyd Reed and Peter Marko for comments and advice that improved the manuscript.

